# Exogenous application of non-mature miRNA-encoded miPEP164c inhibits proanthocyanidin synthesis and stimulates anthocyanin accumulation in grape berry cells

**DOI:** 10.1101/2021.03.04.433997

**Authors:** Mariana Vale, Jéssica Rodrigues, Hélder Badim, Hernâni Gerós, Artur Conde

## Abstract

Secondary metabolic pathways in grape berries are tightly regulated by an array of molecular mechanisms, including microRNA-mediated post-transcriptional regulation. As recently discovered, before being processed into mature miRNAs, the primary transcripts of miRNAs (pri-miRNAs) can encode for small miRNA-encoded peptides (micropeptides - miPEPs) that ultimately led to an accentuated downregulation of the respective miRNA-targeted genes. Although few studies about miPEPs are available, the discovery of miPEPs reveals a new layer of gene regulation at the post-transcriptional level and may present a key advantage in agronomy.

Here, we identified a miPEP encoded in non-mature miR164c putatively targeting grapevine’s transcription factor VvMYBPA1 (miPEP164c/miPEP-MYBPA1), a positive regulator of key genes in the proanthocyanidin-biosynthetic pathway, one that competes directly for substrate with the anthocyanin-biosynthetic pathway.

Thus, the objective of this work was to test the hypothesis that the exogenous application of miPEP164c (miPEP-MYBPA1) can modulate the secondary metabolism of grape berry cells by inhibiting PA biosynthetic pathway while simultaneously stimulating anthocyanin synthesis.

The exogenous application of miPEP164c to suspension-cultured cells from grape berry (cv. Gamay) enhanced the transcription of its corresponding pri-miR164c, thus leading to a more pronounced post-transcriptional silencing of its target VvMYBPA1. This led to a significant inhibition of the proanthocyanidin pathway, mostly via inhibition of *leucoanthocyanidin reductase* and *anthocyanidin reductase* enzymatic activities and Vv*LAR1* downregulation. In parallel, the anthocyanin-biosynthetic route was stimulated. Anthocyanin content was 31 % higher in miPEP164c-treated cells, in agreement with the higher activity of VvUFGT and the corresponding *VvUFGT1* transcripts.

## Introduction

Although grapevines are well adapted to semi-arid climate, the increasingly more frequent combined effect of drought, high air temperature and high evaporative demand has a negative impact in grapevine yield (Chaves et al., 2010) and, if severe, also in berry quality (Teixeira et al., 2013). Therefore, berry and wine quality depend strongly on the grapevine adaptability to drought, heat and light/UV intensity. This abiotic stressors particularly impact highly-regulated molecular mechanisms underlying the synthesis of several quality-related compounds, such as anthocyanins, proanthocyanidins (PAs), flavanols and flavonols (Downey et al., 2006; Teixeira et al., 2013).

Besides being the key component in red wine color, anthocyanins have several health-related properties such as anti-inflammatory and antioxidant capacity, protecting DNA mainly of damage induced by free radicals or reactive oxygen species due to photo-oxidative stress (Gould, 2004). Anthocyanin and proanthocyanidin (condensed tannins) biosynthetic pathways share two common precursors, leucoanthocyanidins and anthocyanidins, and as a result they are in constant competition for carbon availability (Li et al., 2016). Proanthocyanidins are composed of several monomers of catechin and epicatechin, both flavan-3-ols that originate in a branch deviation of the general flavonoid pathway. Catechin synthesis is catalyzed by *leucoanthocyanidin reductase* (LAR), an enzyme that uses leucoanthocyanidins as substrate. However, leucoanthocyanidins can also be catalyzed *by leucoanthocyanidin oxygenase* (LDOX), continuing the flavonoid pathway and resulting in the formation of anthocyanidins, a substrate of both *UDP-glucose flavonoid 3-O-glucosyltransferase* (UFGT), in the synthesis of anthocyanins, and *anthocyanidin reductase* (ANR), in the synthesis of epicatechin, another building block of proanthocyanidins (Gagné, Lacampagne, Claisse, & Gény, 2009). In grapevine, many transcription factors belonging to the R2R3-MYB family are involved in the regulation of flavonoid synthesis by inducing or silencing key biosynthetic genes along the flavonoid pathway (Matus et al., 2009) (Deluc, 2006). The transcription factors VvMYB5a and VvMYB5b are already described as positive regulators of the flavonoid pathway, inducing an upregulation of late-stage berry-associated genes such as *VvCHI* (chalcone isomerase), *VvF3’5* (flavonoid 3′,5′-hydroxylase), *VvDFR* (dihydroflavonol 4-reductase), *VvLDOX*, *VvANR* and *VvLAR1* leading to the synthesis of flavonols, anthocyanidins and proanthocyanidins (Pérez-Díaz et al., 2016) (Cavallini et al., 2014). *VvMYBPA1,* expressed during flowering and early berry development, is a positive regulator of proanthocyanidin (PA) synthesis, by upregulating *VvLDOX, VvANR* and *VvLAR1* genes (Cavallini et al., 2015) (Bogs et al., 2007), thus limiting the progress of the anthocyanin-biosynthetic route.

Regulation of the flavonoid pathway can also be coordinated at the post-transcriptional level by several microRNAs (miRNAs) (Xie et al., 2010) that negatively regulate the expression of their target genes, either by promoting degradation of such target messenger RNAs (mRNAs) or by leading to inhibition of targeted mRNA translation (Pantaleo et al., 2010). MicroRNAs are initially transcribed as much larger primary transcripts (pri-miRNAs) and only become mature miRNA after a maturation processes occurs in the cytosol (Xie et al., 2010). Like any other protein-coding gene, miRNAs genes are transcribed by RNA polymerase II originating the primary transcript of miRNA (pri-miRNA) that consists of a few hundred bases, a 5’cap and 3’ploy-A tail and the characteristic stem-loop structure where the miRNA sequence is inserted, and which is recognized by members of the Dicer-like1 family enzymes. This enzyme cleaves the 5’cap and 3’ poly-A tail of the primary transcript, transforming it in a precursor miRNA (pre-miRNA). DCL1 also carries out the subsequent cleavage of pre-miRNA to release the miRNA:miRNA* duplex which is then translocated to the nucleus by HASTY transporter where the correct miRNA strand is incorporated in a ribonuclear particle to form the RISC complex, the machinery that mediates miRNA-mediated gene silencing (Budak & Akpinar, 2015).

In a groundbreaking finding, it was discovered that, before being processed into mature miRNAs, some pri-miRNAs contain small open reading frames (ORF) that could encode for small regulatory peptides called miRNA-encoded peptides (miPEPs) (Lauressergues et al., 2015). The mechanism of action of miPEPs is by enhancing the transcription and accumulation of the corresponding pri-miRNA, in a sort of positive feedback loop, that subsequently results in accentuated downregulation of the respective miRNA-targeted genes. (Couzigou et al, 2015). For instance, the overexpression of miPEP171b in *Medicago truncatula* led to the increased accumulation of endogenous miR171b (involved in the formation of lateral roots), which resulted in significant changes in root development (Couzigou et al, 2016). Moreover, in soybean (*Glycine max*), it was demonstrated for the first time that the exogenous application of well-chosen, synthetic miPEP172c had a positive impact in nodule formation, by inducing the overexpression of pri-miR172c, whose correspondent miR172c accumulation results in an increase in nodule formation and consequent improvement of N fixation (Couzigou et al., 2016)

More recently, Sharma et al. reported a miPEP in Arabidopsis, miPEP858, by screening the 1000 bp region upstream of pre-miR858 for small ORFs. They found miPEP858 was able to modulate the expression of targets gene involved in plant growth and development and also on the phenylpropanoid pathway, by inducing the expression of pri-miR858 (Sharma & Kamal Badola, 2019).

Although screening for small ORFs, either in the precursor sequence of miRNA or in the region upstream of such precursor miRNA seems to be the mainstream method for finding putative miPEPs when you already have a miRNA or targeted gene in mind, others were also successful in screening for miPEPs using alternative molecular approaches such as homology based computational analysis using expressed sequence tags (ESTs) of a certain species genome by blasting it against miRNA sequences already described, to find homologous of miRNAs and then repeat the same methodologies of miRNA target prediction and pre-miRNA screening for small ORFs (Ram, Mukherjee, & Pandey, 2019).

Finally, in 2020, Chen and colleagues reported a miPEP in grapevine, miPEP171d1 which originated from miR171, a conserved miRNA within different plant species associated with root organ development and capable of promoting adventitious root formation and therefore able to overcome challenges in clonal propagation of *Vitis vinifera*, namely the difficulty in rooting in the cutting and layering process of grapevine (Chen et al., 2020). They screened for ORFs in the 500 bp region upstream of the pre-miR171d and found three small ORFs which by transient expression and promoter activity assays found that the peptide encoded by the first ORF was able to increase the expression of vvi-MIR171d, thus named miPEP171d.

Although few studies about miPEPs are available, the discovery of miPEPs reveals a new layer of gene regulation at the post-transcriptional level and may present a key advantage in agronomy.

Taking these groundbreaking discoveries as basis, the objective of this work was to test the hypothesis that the exogenous application of a newly-identified putative grapevine miPEP by our group (miPEP164c – miPEP-MYBPA1) can modulate the secondary metabolism of grape berry cells by inhibiting PA biosynthetic pathway while simultaneously stimulating anthocyanin synthesis. The micropeptide miPEP164c is putatively targeting MYBPA1, as predicted *in silico*, a gene encoding for a transcription factor that acts as a positive regulator of proanthocyanidin synthesis by activating the expression of VvLAR and VvANR, the enzymes responsible for catechin and epicatechin synthesis, the building blocks of proanthocyanidins (Bogs et al., 2007). For that, a wide array of molecular biology and classic biochemistry approaches were combined to better assess the impact of miPEP164c exogenous treatments on the transcription of key genes involved in secondary metabolic pathways, on the biochemical activity of the corresponding key enzymes, and on the final concentration of secondary metabolites such as anthocyanins, proanthocyanidins and total phenolics.

## Material and Methods

### In silico analyses

A series of *in silico* analyses to identify potential miPEPs in grapevine by combining several bioinformatic tools and databases such as the bioinformatic tool psRNATarget Finder (Dai et al., 2018), a plant small regulatory RNA target predictor, with the aid of GenBank, was used to retrieve information on which, how and where (in the target RNA) miRNAs putatively regulated key genes directly or indirectly involved in the flavonoid biosynthetic pathway. The identified mature miRNAs of those targets of interest were then screened in miRBase (microRNA database) (Kozomara & Griffiths-jones, 2014) for their stem-loop sequences or pri-miRNA sequence, the non-mature sequence of the regulatory miRNAs possibly harboring small open reading frames (ORFs) corresponding to regulatory miPEPs. Finally, the obtained stem-lop sequences were then ran in a bioinformatic ORF finder tool that recognizes in the introduced sequence all possible ORFs that could translate into a small peptide, by defining several parameters based on the few miPEPs so far identified in the literature (Lauressergues et al., 2015).

### Solubilization of miPEPs

Following *in silico* identification, the miPEP sequence was ordered from Smart Bioscience as 1 mg aliquot. Solubilization of the micropeptide was conducted as recommended by Smart-Bioscience Peptide Solubility Guidelines (https://www.smart-bioscience.com/support/solubility/). miPEP164c, putatively negatively regulating *VvMYBPA1*, was solubilized in 200 μL of acetic acid (10%) and 800 μL of DMSO to a final concentration of 1 mg/mL. A solution of 200 μL of acetic acid (10%) and 800 μL of DMSO was used as control in the exogenous treatments.

### Biological material

Grape berry cell suspensions of the cultivar Gamay Freaux cv. were maintained in Gamborg B5 medium in 250 mL flasks at 25 °C with constant agitation on a rotator shaker at 100 rpm and under 16h/8 h photoperiod. The culture medium composition was as follows: 3 g/L Gamborg B5 salt mixture and vitamins; 30 g/L sucrose (3% m/v); 250 mg/L casein enzymatic hydrolysate; 0.1 mg/L α-napthaleneacetic acid (NAA); 0.2 mg/L Kinetin, and a final pH of 5.7. The suspension-cultured cells were allowed to grow for 10 d, until the exponential phase, when they were subcultured by transferring 10 mL of cells to 30 mL of fresh medium.

### Exogenous addition of micropeptides to Gamay grape cells

For each assay, immediately after sub-cultivation, 1 μM of miPEP164c was exogenously added to the cell cultures, in a volume that represented no more than 0.15 % (v/v) of the total volume of the cell suspension. All cell suspensions, including control cells (treated with the same volume of control solution) were cultivated for 10 d with constant agitation and a 16h/8 h photoperiod. Cells were then collected, filtered and immediately frozen with liquid nitrogen and stored at −80 °C. A part of cells of each experimental condition was lyophilized.

### Quantification of anthocyanins

Anthocyanins were extracted from 100 mg of grape berry cells from each experimental condition. After adding 1 mL of 100% methanol, the suspensions were vigorously shaken for 30 min, following centrifugation at 18.000*xg* for 20 min. The supernatants were collected and 200 μL of each supernatant was mixed with 1.8 mL of a solution of 25 mM KCl (pH = 1.0) and absorbance was measured at 520 nm and 700 nm. Total anthocyanin quantification was calculated in relation to cyanidin-3-glucoside equivalents, as follows:

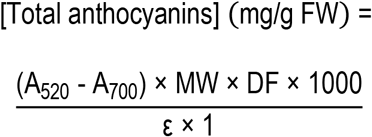

where MW is the molecular weight of cyanidin-3-glucoside (449.2 g mol^−1^), DF is the dilution factor and ε is the molar extinction coefficient of cyanidin-3-glucoside (26900 M^−1^ cm^−1^).

### Quantification of proanthocyanidins

Proanthocyanidin content was determined using an adapted colorimetric vanillin-HCl assay described by Broadhurst & Jones (1978). To extract proanthocyanidins, 1 mL of 100% methanol was added to 5 mg of lyophilized grape berry cells and vigorously shaken for 30 min followed by centrifugation at 18000*xg* for 15 min. Supernatants were collected and diluted in a 1:1 ratio with methanol to final volume of 500 μL. The methanolic extracts were added to clean assay tubes wrapped with aluminum foil. Then, 3 mL of a solution of 4% (m/v) vanillic acid freshly prepared in methanol was added and mixed very gently. Finally, 1.5 mL of concentrated hydrochloric acid was added to each reaction tube and mixed very gently. The reactions were allowed to stand for 6 min and the absorbance of the samples was measured spectrophotometrically at 500 nm. To discard absorbance interference caused by anthocyanin presence in the samples, control reactions for each condition were prepared with 3 mL of methanol instead of vanillic acid and the absorbance measured at 500 nm was discounted from the absorbance of reaction mixtures with vanillic acid. An epigallocatechin gallate (EGCG) standard curve, with concentrations ranging from 10 to 200 μg employing the same method was always prepared for each quantification of proanthocyanidin content.

### Protein extraction

Protein extraction was conducted as described in Conde et al. (2016). Lyophilized grape berry cells were mixed with extraction buffer in approximately 1:1 (v/v) powder: buffer ratio. Protein extraction buffer contained 50 mM Tris-HCl, pH 8.9, 5 mM MgCl_2_, 1 mM EDTA, 1 mM PMSF, 5 mM DTT and 0.1 % (v/v) Triton X-100. Homogenates were thoroughly vortexed and centrifuged at 18000*xg* for 15 min at 4 °C. Supernatants were maintained on ice and used for all enzymatic assays. Total protein concentrations of the extracts were determined by the method of Bradford (Bradford, 1976) using bovine serum albumin as a standard.

### Enzymatic activity assays

The biochemical activity of UDP-Glucose:flavonoid 3-O Glucosyltransferase (UFGT) was determined as described by Conde et al. (2016) with some adaptations. The assay mixture contained 385 μL of 100 mM Tris-HCl reaction buffer (pH 8), 100 μL of enzyme extract, 10 μL of 50 mM UDP-glucose and the reaction was initiated with 5 μL of 100 mM quercetin as substrate for the enzyme activity (saturating concentration) to a final reaction volume of 500 μL. Each mixture was incubated for 30 min in the dark with gentle shaking. After incubation, dilutions were prepared with 100 μL of each assay mixture and 900 μL of Tris-HCl reaction buffer and absorbance was read at 350 nm immediately after (t=0) and 30 min later (t=30) to follow the production of quercetin 3-glucoside (ε = 21877 M^−1^ cm^−1^).

Leucoanthocyanidin reductase (LAR) enzymatic activity was measured by spectrophotometrically monitoring the conversion of dihydroquercetin to (+)-catechin following the method of Gagné et al. (2009) with some adaptions. The assay mixture contained 1,7 mL of Tris-HCl buffer (0.1 M, pH 7.5), 300 μL of protein extract, 2 μL of NADPH (100 mM) and the reaction was initiated by adding 1 μL of dihydroquercetin (10 mg mL^−1^ in DMSO). The production of (+)-catechin (ε = 10233 M^−1^ cm^−1^) was followed at 280 nm for 30 min.

The biochemical activity of anthocyanidin reductase (ANR) was determined as described by Zhang et al. (2012) with some adaptations. The assay mixture contained 1,5 mL of PBS buffer (0.1 M, pH 6.5), 60 μL of enzyme extract, 40 μL of ascorbic acid (20 mM), 50 μL of cyanidin chloride (2 mM) and the reaction was initiated by adding 75 μL of NADPH (20 mM) followed by a 1/10 dilution with PBS reaction buffer for proper absorbance measure. The enzyme activity was monitored by measuring the rate of NADPH (ε = 6,22 mM^−1^ cm^−1^) oxidation at 340 nm for 20 min, at 45°C.

### RNA extraction and cDNA synthesis

Total RNA extraction was performed according to Reid et al. (2006) in combination with purification steps from the GRS Total Plant RNA extraction kit. After treatment with DNase I (Qiagen), cDNA was synthesized from 1 μg of total RNA using Maxima first strand cDNA synthesis kit from Thermo Scientific, following the manufacturer’s instructions. RNA concentration and purity were determined using Nanodrop and its integrity assessed in a 1% agarose gel stained with SYBR Safe (InvitrogenTM, Life Technologies).

### Transcriptional analyses by real-time qPCR

Quantitative real-time PCR was performed with QuantiTect SYBR Green PCR Kit (Qiagen) in a CFX96 Real-Time Detection System (Bio-Rad), using 1 μL of cDNA in a final reaction volume of 10 μL per well. Specific primer pairs used for each target gene arelisted in Table 1. Melting curve analysis was performed for specific gene amplification confirmation. As reference genes, *VvACT1* (actin) and *VvGAPDH* (glyceraldehyde-3-phosphate dehydrogenase) were selected, as these genes were proven to be very stable and ideal for qPCR normalization purposes in grapevine (Reid et al., 2006). For all experimental conditions tested, two independent runs with triplicates were performed. The expression values were normalized by the average of the expression of the reference genes as described by Pfaffl (2001) and analyzed using the software Bio-Rad CFX Manager (Bio-Rad).

**Table 1.**
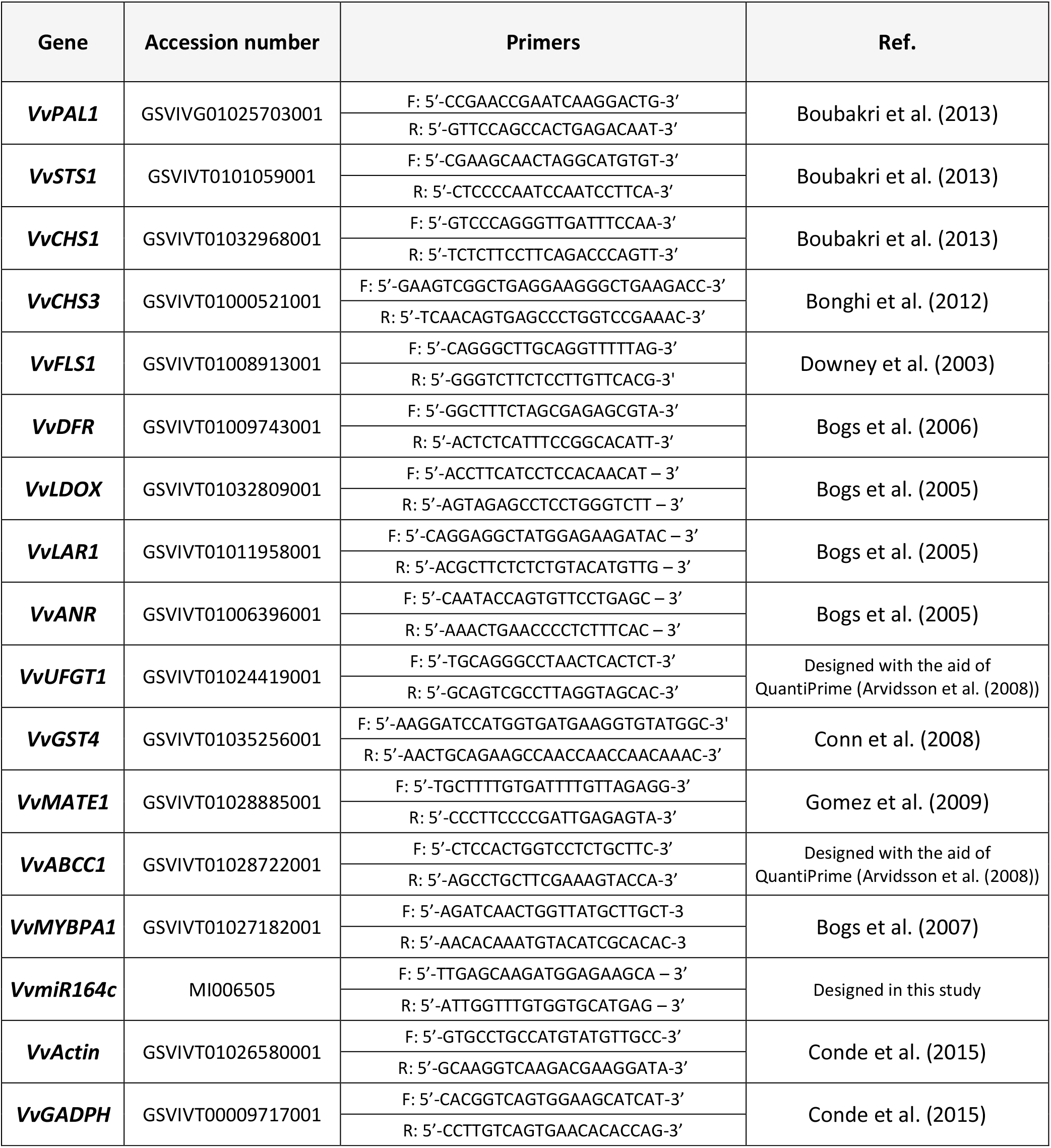
Primers forward (F) and reverse (R) used for gene expression analysis by qPCR.

**Table 2.**
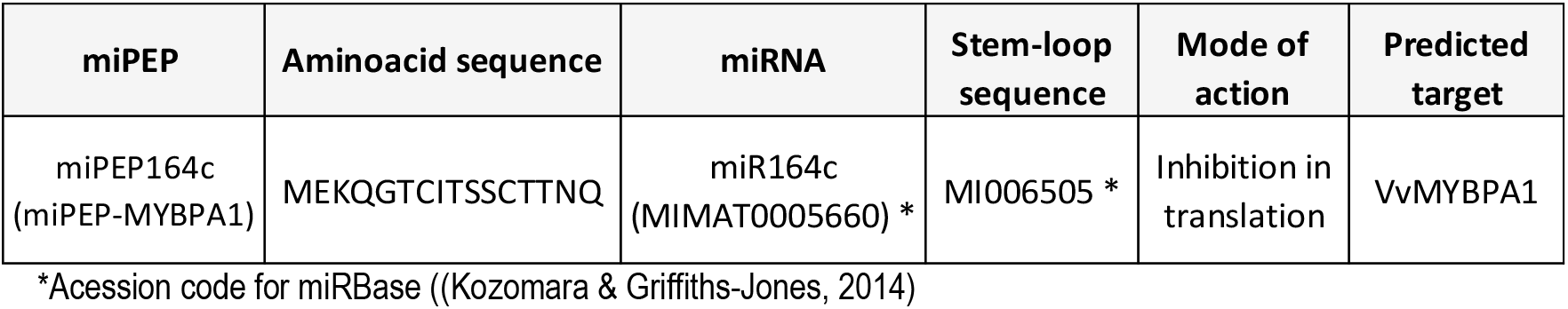
Detailed information about the micropeptide identified by an *in silico* analysis and selected for this study and its corresponding mature miRNA and mode of action

### Statistical analyses

The results were statistically analyzed by Student’s t-test using Prism vs. 6 (GraphPad Software, Inc.). For each condition, statistical differences between mean values are marked with asterisks.

## Results

### Identification and *in silico* analysis of the candidate grapevine micropeptide miPEP164c

An *in silico* analysis for micropeptide screening led to the selection of miPEP164c a candidate miPEP with putative regulatory function in grape berry flavonoid biosynthesis metabolic pathway, particularly in the branch of proanthocyanidin synthesis. miR164c was predicted *in silico* to post-transcriptionally inhibit grapevine transcription factor *VvMYBPA1*, involved in the activation of flavonoid synthesis, specifically of proanthocyanidin synthesis (via LAR1, LAR2 and ANR activation). Relevant information obtained by the *in silico* analysis regarding the miPEP selected for this study, including its aminoacidic sequence, attributed name and respective mature miRNA name and miRbase accession number, as well as that of its precursor miRNA (pre-miRNA), is detailed in Supplementary Table 2.

### Effect of miPEP164c exogenous application on the abundance of miR164c and its putative target transcription factor VvMYBPA1

To confirm if miPEP164c exogenous application is indeed activating the accumulation of its *in silico* predicted miRNA (miR164c), gene expression analysis by real-time qPCR of the non-mature pre-miR164c was performed on cells treated with miPEP164c. As shown in Figure 1A, 10 days after treatment, the transcript levels of pre-miR164c were stimulated (3.5-fold increase) by miPEP164c exogenous application. A slight, non-significant increase in the transcript levels of *VvMYBPA1* was also observed after 10 days of miPEP164c treatment (Fig. 1B).

**Figure 1.**
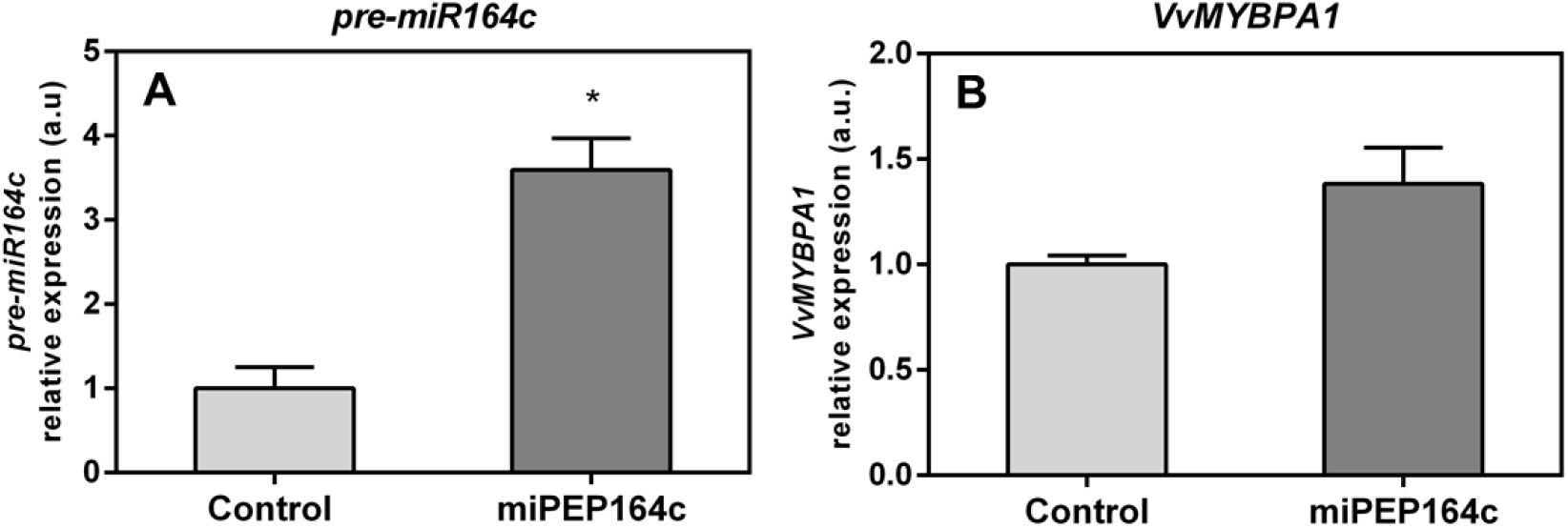
Steady-state transcript levels of *pre-miR164c* (A) and *VvMYBPA1* (B) in suspension-cultured grape berry cells (cv. Gamay) 10 d after elicitation with 1 μM miPEP164c. Gene expression analysis, by real-time qPCR was normalized with the expression of reference gene *VvACT1* and *VvGAPDH*. Values are the mean ± SEM. Asterisks indicate statistical significance (Student’s t-test; * P < 0.05).

### Effect of miPEP164c exogenous application on grape berry key secondary metabolites

Spectrophotometric quantifications of grape berry cells secondary metabolites revealed a significant increase of 31% in anthocyanin content (Fig. 2A) while total proanthocyanidins significantly decreased by 26% (Fig. 2B), after 10 days of treatment with 1 μM miPEP164c.

**Figure 2.**
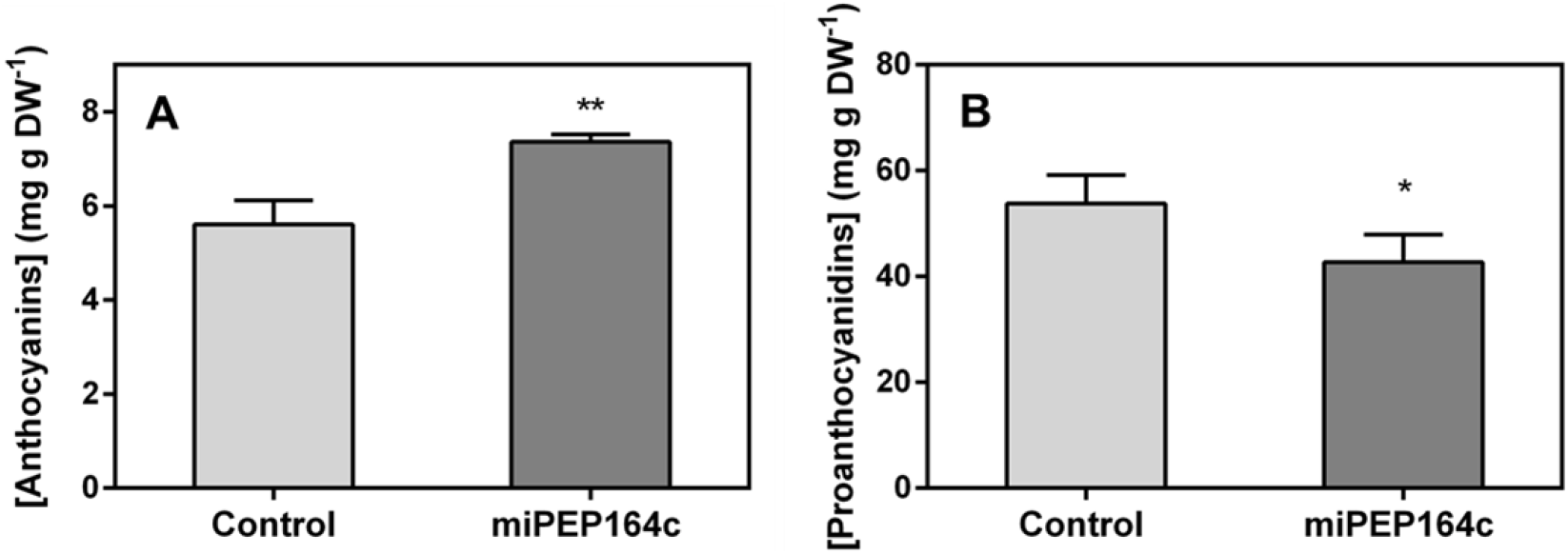
Effect of the exogenous application miPEP164c (1 μM) on total anthocyanin content (A) and on total proanthocyanidin content (B) in suspension-cultured grape berry cells (cv. Gamay) 10 d after elicitation with 1 μM miPEP164c. Anthocyanin concentration is represented as mg of cyanidin 3-glucoside (C-3-G) equivalents per g of fresh weight (FW). Asterisks indicates statistical significance (Student’s t-test; *P<0.05 ** P<0.01).

### Transcriptional and biochemical changes induced by miPEP164c on the proanthocyanidin-synthesizing branch

Analysis by real-time qPCR showed that *VvLAR1* expression was reduced by 20% in Gamay cells elicited with 1 μM miPEP164c compared to control cells (Fig. 3B). In agreement with the observed decrease in *VvLAR1* expression levels under miPEP164c treatment, the specific activity of LAR was 3-fold reduced (Fig. 3A).

**Figure 3.**
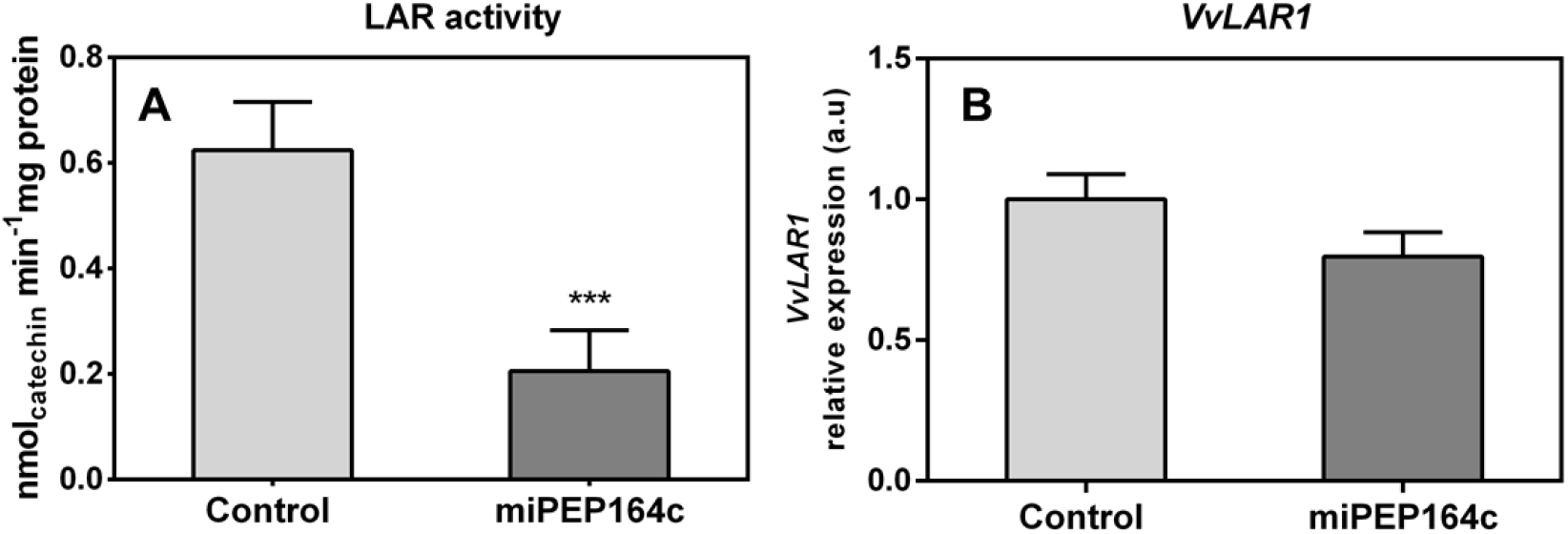
Effect on the specific activity of LAR (A) and steady-state transcript levels of *VvLAR1* (B) in suspension-cultured grape berry cells (cv. Gamay) 10 d after elicitation with 1 μM miPEP164c. Gene expression analysis, by real-time qPCR was normalized with the expression of reference gene *VvACT1* and *VvGAPDH*. Values are the mean ± SEM. Asterisks indicate statistical significance (Student’s t-test; *** P < 0.001). LAR biochemical activity represented as the *V*_max_ in grape berry cells under miPEP164c treatment. Values are the mean ± SEM. Asterisks indicates statistical significance (Student’s t-test; *** P < 0.001)

The specific activity of ANR also decreased (by 27%) in Gamay cells elicited with 1 μM miPEP164c (Fig. 4A), but *VvANR* expression was not repressed, instead a 18% increase of the steady-state transcript levels was observed, although statistically not significant (Fig. 4B)

**Figure 4.**
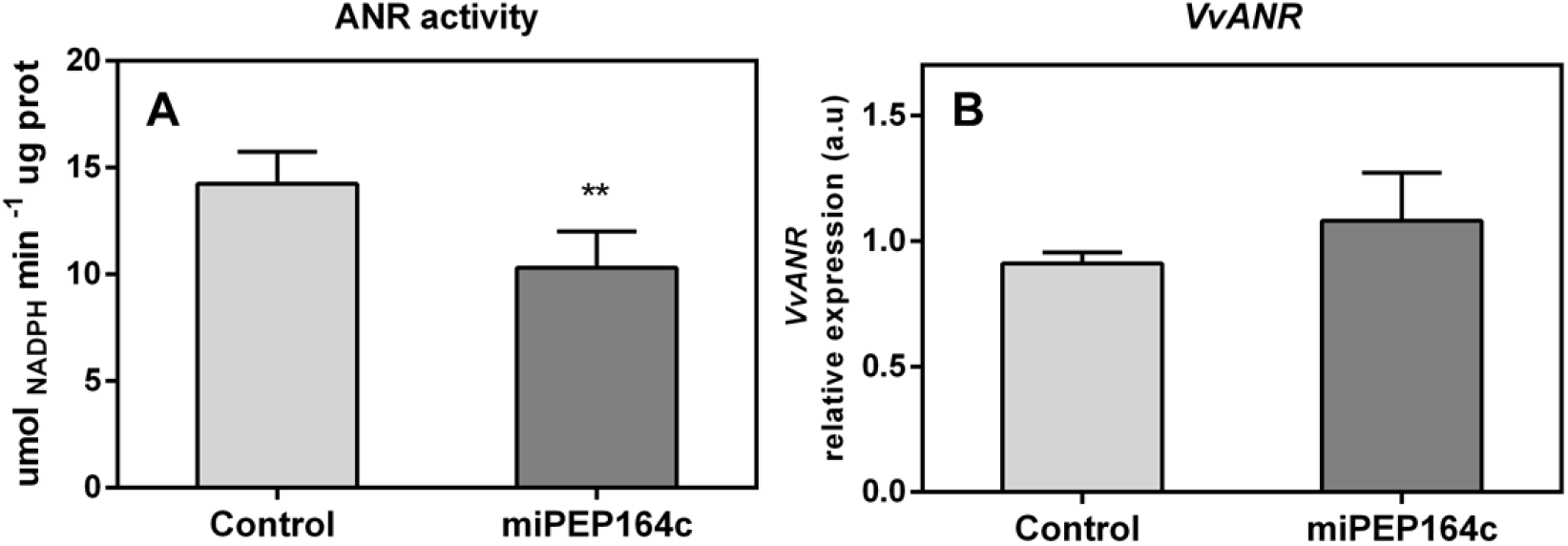
Effect on the specific activity of ANR (A) and steady-state transcript levels of *VvANR* (B) in suspension-cultured grape berry cells (cv. Gamay) 10 d after elicitation with 1 μM miPEP164c. Gene expression analysis, by real-time qPCR was normalized with the expression of reference gene *VvACT1* and *VvGAPDH*. Values are the mean ± SEM. ANR biochemical activity, represented as the *V*_max_ in grape berry cells under miPEP164c treatment. Values are the mean ± SEM. Asterisks indicates statistical significance (Student’s t-test; ** P < 0.01)

### Transcriptional and biochemical changes induced by miPEP164c on the anthocyanin-synthesizing branch

The expression of *VvUFGT1* was strongly stimulated by miPEP164c application, reflected by a 4-fold increase in the expression levels in grape berry cells under this treatment (Fig. 5B), which corroborates with a significant increase in the total concentration of anthocyanins observed previously (Fig. 2A). In agreement with *VvUFGT1* transcripts increase, the biochemical activity of UFGT was 3.2-fold higher in miPEP164c treated cells, reaching a *V*_max_ of 3.4 μmol h^−1^ mg protein^−1^ (Fig. 5A).

**Figure 5.**
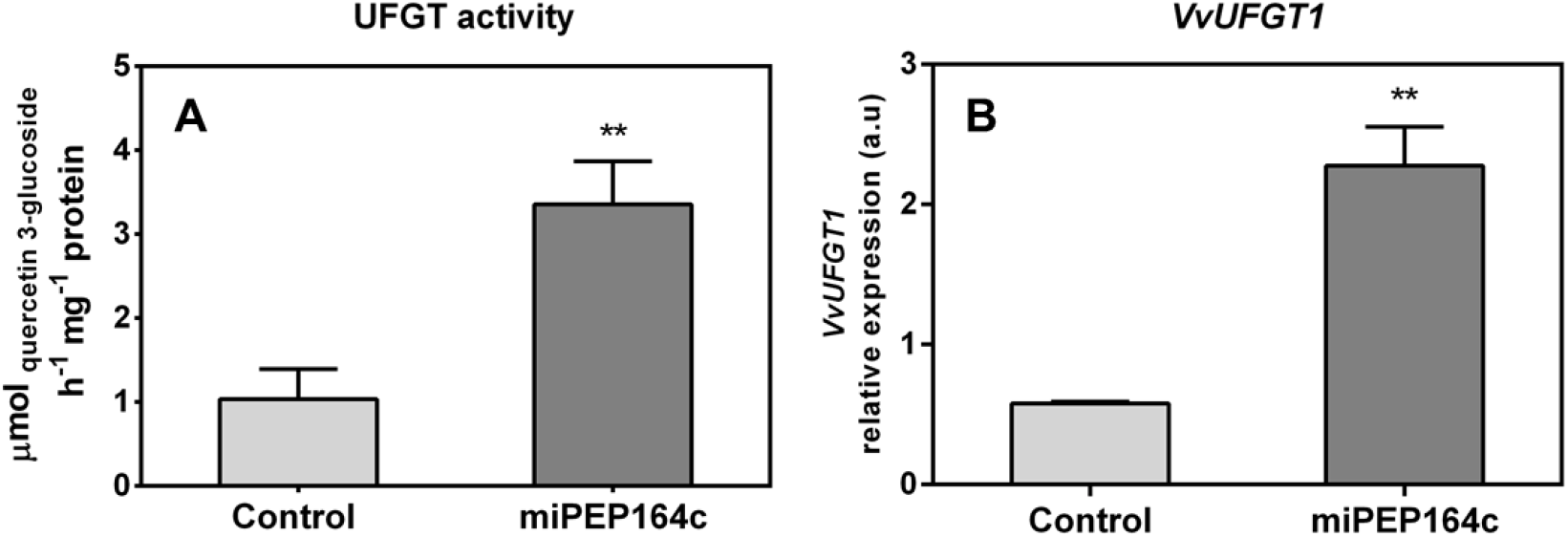
Effect on the specific activity of UFGT (A) and steady-state transcript levels of *VvUFGT1* (B) in suspension-cultured grape berry cells (cv. Gamay) 10 d after elicitation with 1 μM miPEP164c. Gene expression analysis, by real-time qPCR was normalized with the expression of reference gene *VvACT1* and *VvGAPDH*. Values are the mean ± SEM. Asterisks indicate statistical significance (Student’s t-test; ** P < 0.01). UFGT biochemical activity, represented as the *V*_max_ in grape berry cells under miPEP164c treatment. Values are the mean ± SEM. Asterisks indicates statistical significance (Student’s t-test; ** P < 0.01)

The expression levels of *VvDFR* were also significantly stimulated, with an increase of 2-fold in grape berry cells 10 days after miPEP164c treatment (Fig. 6A). *VvLDOX* was also significantly stimulated under this treatment, increasing its expression levels by 42% when compared to control cells (Fig. 6B).

**Figure 6.**
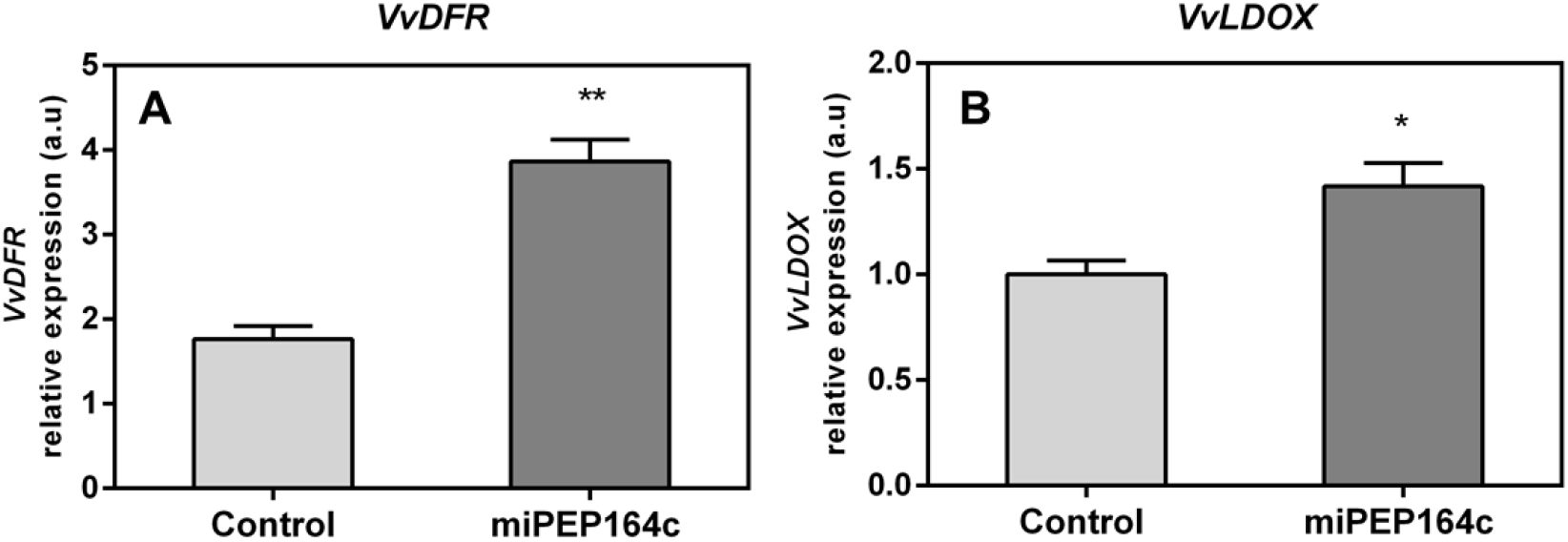
Steady-state transcript levels of *VvDFR* (A) and *VvLDOX* (B) in suspension-cultured grape berry cells (cv. Gamay) 10 d after elicitation with 1 μM miPEP164c. Gene expression analysis, by real-time qPCR was normalized with the expression of reference gene *VvACT1* and *VvGAPDH*. Values are the mean ± SEM. Asterisks indicate statistical significance (Student’s t-test; * P < 0.05, ** P < 0.01).

Transcriptional analysis showed that the expression of *VvGST4* under miPEP164c treatment also increase 2-fold (Fig 7A). Similarly, the transcript levels of *VvMATE1* increased by 55% (Fig 7B), while the expression of *VvABCC1* did not seem to be affected by treatment with this micropeptide (Fig. 7C). These genes encode transporters that accumulate anthocyanins in the vacuole.

**Figure 7.**
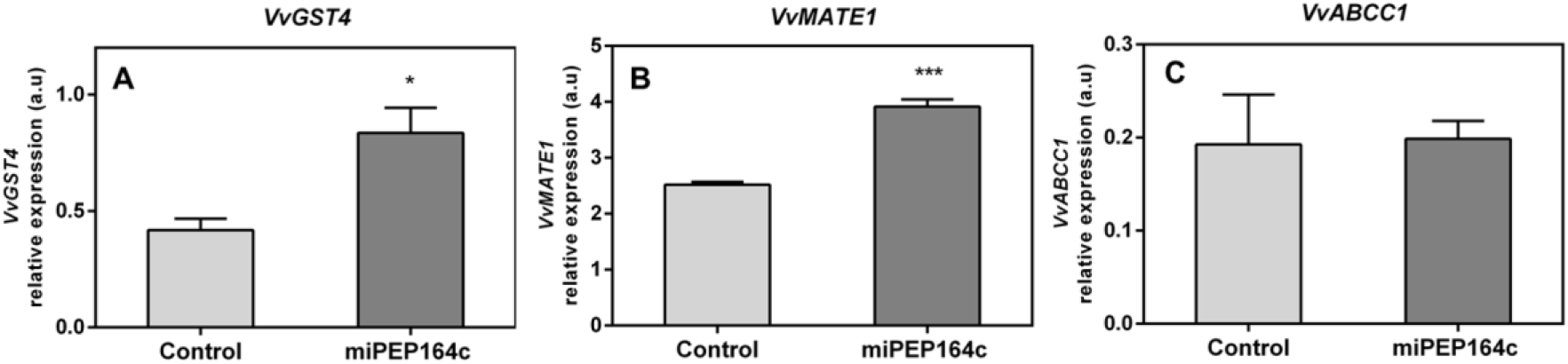
Steady-state transcript levels of *VvGST4* (A), *VvMATE1* (B) and *VvABCC1* (C) in suspension-cultured grape berry cells (cv. Gamay) 10 d after elicitation with 1 μM miPEP164c. Gene expression analysis, by real-time qPCR was normalized with the expression of reference gene *VvACT1* and *VvGAPDH*. Values are the mean ± SEM. Asterisks indicate statistical significance (Student’s t-test; * P < 0.05, *** P < 0.001).

## Discussion

Grape berry secondary metabolism generates, by a cascade of reactions scattered through different branches of the phenylpropanoid pathway, a wide range of bioactive compounds with key roles in plant defense responses and with several health-related benefits to humans, making them metabolites of interest for many industries (Teixeira et al., 2013). Therefore, the search for new strategies to modulate these complex pathways in the hopes of either minimizing the effects of several stress factors in the composition and quality of grape secondary metabolites or to increase the synthesis and accumulation of bioactive metabolites of interest such as antioxidant compounds like anthocyanins, is a research line of great importance, not only to the viticulture industry but that can also be applied in several industries of health-promoting products (Zhang et al., 2015). In the present study, we sought to validate a new and promising strategy to modulate the secondary metabolism of Gamay grape berry cells by testing a synthetic miPEP, putatively enhancing the transcription and accumulation of miR164c and ultimately promoting a more pronounced silencing of its predicted target. Because this transcription factor is involved in the molecular activation of key genes in the proanthocyanidin pathway, ultimately, we wanted to evaluate if a miPEP-based treatment could regulate grape berry secondary metabolism, by activating miRNA-mediated post-transcription silencing mechanisms of specific targets. Results obtained were very promising as this treatment could represent an innovative and easy-to-apply strategy to modulate the synthesis of more quality-related compounds, resulting in crops with added-value characteristics, without the need for more drastic, time consuming and more expensive strategies, as genetic transformation of crops.

### Elicitation of Gamay cells with miPEP164c induces accumulation of miR164c and consequent miR164c-mediated inhibition of proanthocyanidin biosynthetic pathway

Overall, results confirmed that the exogenous application of miPEP164c is indeed enhancing the accumulation of miR164c which ultimately resulted in a more pronounced post-transcriptional silencing of transcription factor VvMYBPA1 and consequently of MYBPA1-activated genes, here observed by a significant downregulation of LAR and ANR specific activity resulting in a significant decrease of 26% of total proanthocyanidin content in cells under miPEP164c treatment.

Gene expression analysis by real-time qPCR confirmed that miPEP164c increased the expression levels of the pre-miR164c in Gamay cells. Thus, a positive loop was established, in which a consequent increased translation into miPEP164c, ultimately results in higher levels of mature miR164c and accentuated negative regulation of the target gene *VvMYBPA1*. *In silico* analyses suggests that mode of action of miR164c is through inhibition of the translation of *VvMYBPA1*, not by cleavage of the target messenger RNA, due to a lack of 100% complementarity between the guide miRNA and the target mRNA (Waterhouse & Hellens, 2015). This goes in agreement with our results showing that the treatment with miPEP164c did not induce any significant changes in the expression levels of *VvMYBPA1*.

Evidence for the involvement of post-transcriptional silencing of *VvMYBPA1* mediated by miPEP164c was obtained when the MYBPA1-activated enzyme *VvLAR* was clearly down-regulated. Both *VvANR* and *VvLAR1*, are key genes leading to the synthesis of proanthocyanidins (Gagné et al., 2009). However, the expression of *VvANR*, encoding for the enzyme that synthesizes epicatechins from anthocyanidins, was not affected, possibly to compensate the decreased activity of VvLAR, in order to ensure a certain amount of monomers for proanthocyanidins biosynthesis. Also, *VvANR* expression may be regulated by several other regulatory proteins, such as *VvMYC1*, a bHLH transcription factor that physically interacts with MYB-like transcription factors like MYBPA1 and MYB5a/b to coordinate the regulation of *VvANR*, and therefore silencing of one regulator may be overcome by another regulatory mechanism (Heppel, 2010).

### Proanthocyanidin synthesis was inhibited by miPEP164c while anthocyanin synthesis was simultaneously increased

The observed significant increase in anthocyanin total content in Gamay cells mediated by the application of miPEP164c corroborates our hypothesis that a miPEP164c-mediated silencing of proanthocyanidin synthesis would divert the carbon flow to the anthocyanin branch, due to the constant competition of both pathways for the same substrates, as reported before (Liao et al., 2015). Gene expression analysis of *VvUFGT1,* that glycosylates anthocyanidins into anthocyanins, revealed a strong upregulation of its expression levels in response to the elicitation with the micropeptide which goes in agreement with the observed increase of the UFGT specific activity that also increased.

In *V. vinifera* two types of anthocyanin tonoplast transporters that accumulate anthocyanins in the vacuole were identified: primary transporters from the ATP-binding cassette (ABC) family, such as the *VvABCC1* who requires the presence of reduced glutathione (GSH) to properly transport anthocyanins, through the tonoplast, into the vacuole (Jiang et al., 2019); and tonoplast secondary transporters like *VvMATE1* (anthoMATE) of the multidrug and toxic extrusion family that use the H^+^ gradient to transport mostly acylated anthocyanins (Gomez et al., 2009). Also crucial for anthocyanin stabilization and transport are the glutathione S-transferases, as the paradigmatic case of grapevine’s *VvGST4*, to promote anthocyanin S-conjugation with reduced glutathione for anthocyanin-stabilization purposes (Conn et al., 2008). Several studies on the role of GSTs in anthocyanin accumulation have described GSTs as escort/carrier proteins, binding anthocyanins to form a GST-anthocyanin complex, protecting them from oxidation and guiding anthocyanins from the cytosolic surface of the ER to the vacuole for proper storage mediated by tonoplast transporters such as *VvMATE1* and *VvABCC1* (Zhao & Dixon, 2010) (Jiang et al., 2019). Our results strongly supported that anthocyanin transport capacity to the vacuole, where they are stored in grape berry cells, was also stimulated by miPEP164c application as the expression of the anthocyanin tonoplast transporter *VvMATE1* and anthocyanin carrier protein *VvGST4*, was upregulated by this micropeptide. It is not understood how plants choose between ATP-hydrolysis-dependent or H+/Na+-gradient dependent mechanisms for transport of native metabolites or xenobiotics. However, it is believed that the conjugation ligands, such as glucose or glutathione, play a key role in the determination of which transport mechanism will be used (Zhao & Dixon, 2010). However, the expression of *VvABCC1* was not affected by miPEP164c contrarily to what would be expected considering the upregulation of *VvGST4* expression. This could be due to the presence of other regulatory proteins affecting the expression of *VvABCC1*, other phenolic substrates that also need to be transported by this mechanism, the majority of anthocyanins might not be in the glycosylated form, which is the preferred form of anthocyanins of this type of transporter, or simply because it is competing with the upregulated *VvMATE1* transporter for anthocyanins (Francisco et al., 2013).

The increased anthocyanin concentration in treated cells may have also resulted from the observed stimulatory effect of the micropeptide on the transcription of several intermediates along the flavonoid pathway (*VvCHS1*) and anthocyanin synthesis pathway (*VvDFR, VvLDOX* and *VvUFGT1*). The observation that the micropeptide also induced a slight decrease in the expression of *VvFLS1* corroborates previous studies in flowers where the flavonol branch is constantly competing with the anthocyanin branch for precursors for the synthesis of white pigments and of red to blue pigments, respectively (Zhang et al., 2017).

## Conclusion

In this study, recurring to a combination of molecular and biochemical approaches, we revealed that miPEP164c exogenous application induced a strong up-regulation of genes involved in anthocyanin synthesis, transport, and accumulation in the vacuole. Additionally, miPEP164c provoked a downregulation of proanthocyanidin synthesis (a pathway that directly competes with the anthocyanin-biosynthetic pathway), due to a decrease in *VvLAR1* expression levels with a corresponding very significant decrease in LAR total biochemical activity as well as a significant decrease in ANR biochemical activity.

This upregulation of the anthocyanin biosynthetic route seems to be an indirect effect of miPEP164c putatively inhibiting transcription factor MYBPA1, a known positive regulator of proanthocyanidin synthesis. Thus, these metabolic alterations triggered by miPEP164c clearly resulted in higher concentration of anthocyanins and lower concentration of proanthocyanidins, due to miR164c-mediated negative regulation of proanthocyanidin-related transcription factor *VvMYBPA1* and, consequently, *VvLAR1 and VvANR*, ultimately leading to proanthocyanidin synthesis inhibition and anthocyanin synthesis stimulation as these pathways directly compete for substrate, in a mechanism illustrated in Figure 8.

**Figure 8.**
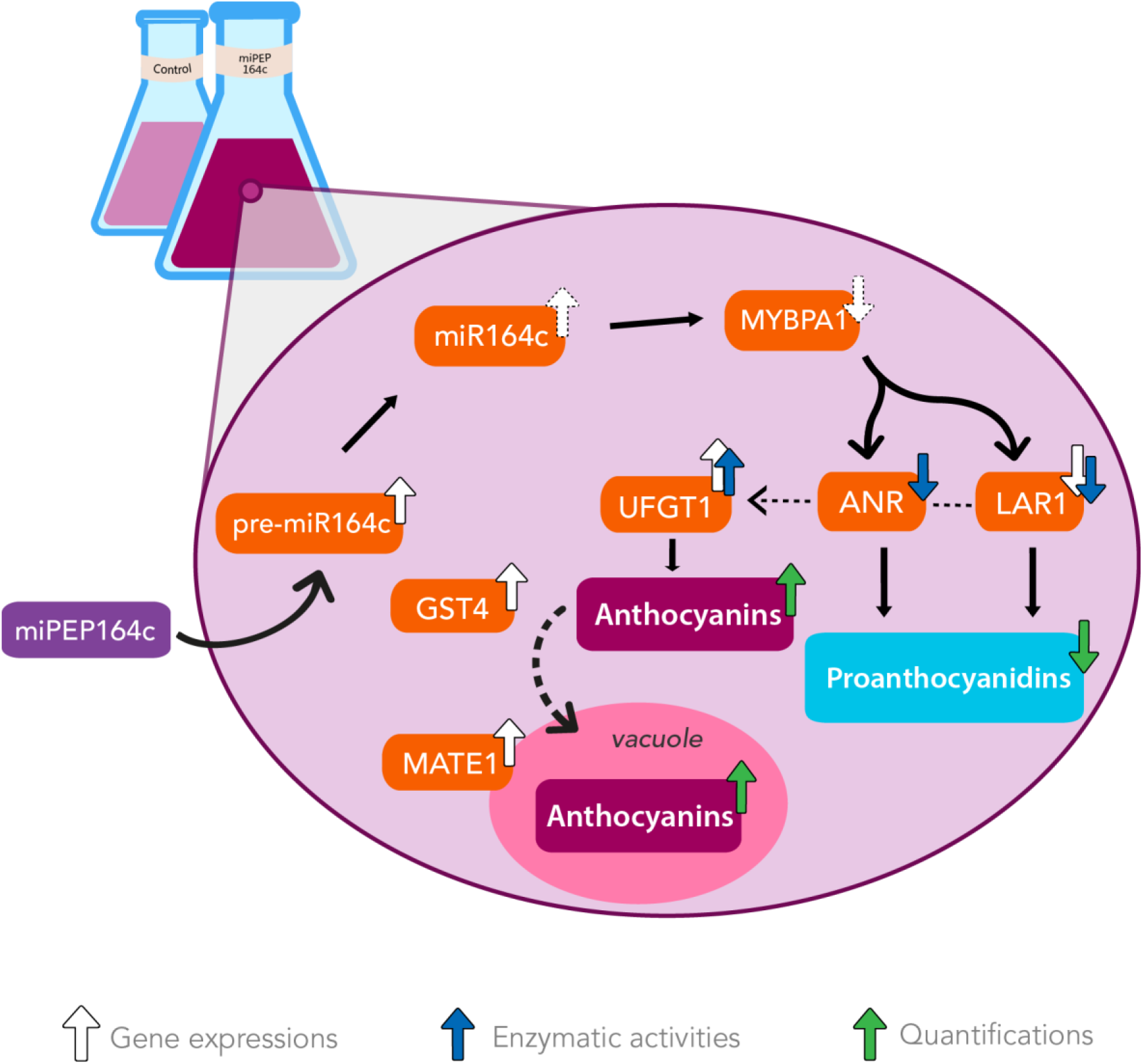
The exogenous application of miPEP164c increases anthocyanin synthesis and accumulation while decreasing proanthocyanidin synthesis by enhancing miR164c-mediated downregulation of PA-synthetic pathway in Gamay grape berry cell suspensions. Addition of miPEP164c provoked an increase in the transcription of pre-miR164c and consequently of its mature form, miR164c, which putatively led to a decrease in the translation of transcription factor MYBPA1. This inhibition in MYBPA1 translation resulted in a downregulation of *VvLAR1* expression and LAR total biochemical activity, as well as ANR total biochemical activity, decreasing the intracellular concentration of proanthocyanidins. This downregulation of the proanthocyanidin pathway indirectly led to the stimulation of the anthocyanin synthesis by increasing *VvUFGT1* expression and UFGT total biochemical activity, as well as vacuolar accumulation, as shown by *VvGST4* and *VvMATE1* overexpression.

Taking together, a miPEP-based treatment is a promising new, cost-efficient strategy to improve plant cell characteristics, with the ability to modulate the secondary metabolism of plants, enhancing the synthesis of quality-related compounds, without the need for genetic engineering of crops. Applied *in planta* it may have the capacity to induce metabolic changes and to be a potential promising technique to improve berry quality or maybe even the resistance of grapevine to biotic stressors such *as Botrytis cinerea* infection, by inducing the production of bioactive compounds that will help plant defense, or even to abiotic stress in the form of extreme UV-radiation due to the anthocyanins role in protecting grape berries against this increasingly more common environmental factor, considering that miPEP164c led to a higher anthocyanin intracellular concentration. Considering this potential, the fact that the exogenous application of miPEPs can specifically modulate plant secondary metabolism may indeed have great agronomical applications.

In a more fundamental research level, further studies are required to fully understand the scope of miPEP-mediated post-transcriptional silencing, as little is known about its mechanisms of action, transport or stabilization in the cytoplasm. Since miPEPs were first discovered, many questions have arisen about their mode of action: how do they interact with the transcriptional machinery to enhance the transcription of the corresponding pri-miRNA? Do all miRNAs have the ability to encode miPEPs? What are the mechanisms that allow synthetic miPEPs to be transported through the cell wall and cell membrane into the nucleus? And how are the primary transcripts of miRNAs able to be translated in the cytoplasm when they are capped and polyadenylated RNA molecules rapidly recognized by the dicing complex?

